## Supplementary Figures for "Exposure to artemisinin at the trophozoite stage increases sexual conversion rates in the malaria parasite *Plasmodium falciparum*"

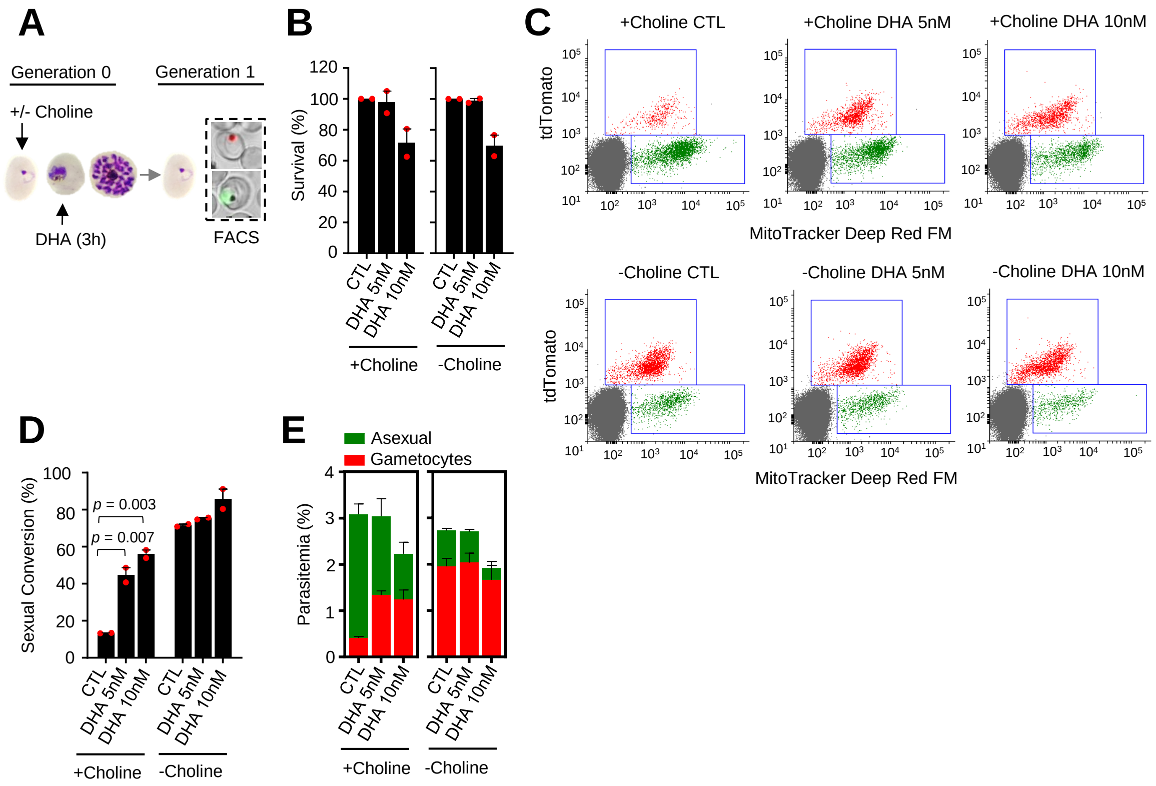


**Supplementary Figure 1. Effect of a dihydroartemisinin (DHA) pulse at the trophozoite stage on sexual conversion, determined using MitoTracker to identify viable parasites. (A)** Schematic representation of the assay. Tightly synchronized cultures of the *NF54-gexp02-Tom* line maintained under non-inducing (+ choline) or inducing (- choline) conditions were exposed to a 3 h DHA pulse at subcurative doses at the trophozoite stage (25-30 hpi). Sexual conversion was measured by flow cytometry (FACS) after reinvasion (~30-35 hpi of the next multiplication cycle). **(B)** Survival rate of cultures exposed to the different drug doses, using total parasitemia of live parasites (asexual + sexual parasites) determined with a mitochondrial membrane potential stain (MitoTracker Deep Red FM). For each choline condition, values are presented relative to the parasitemia in the control cultures (no drug). **(C)** Representative MitoTracker Deep Red FM vs TdTomato (marks gametocytes) flow cytometry dot plots. **(D)** Sexual conversion rates determined by flow cytometry, calculated using MitoTracker-positive cells only. The *p* value is indicated only for treatment vs control (no drug) significant differences ($p<0.05$). **(E)** Distribution of absolute parasitemia of asexual and sexual parasites. In all panels, data are presented as the average and s.e.m. of 2 independent biological replicates.

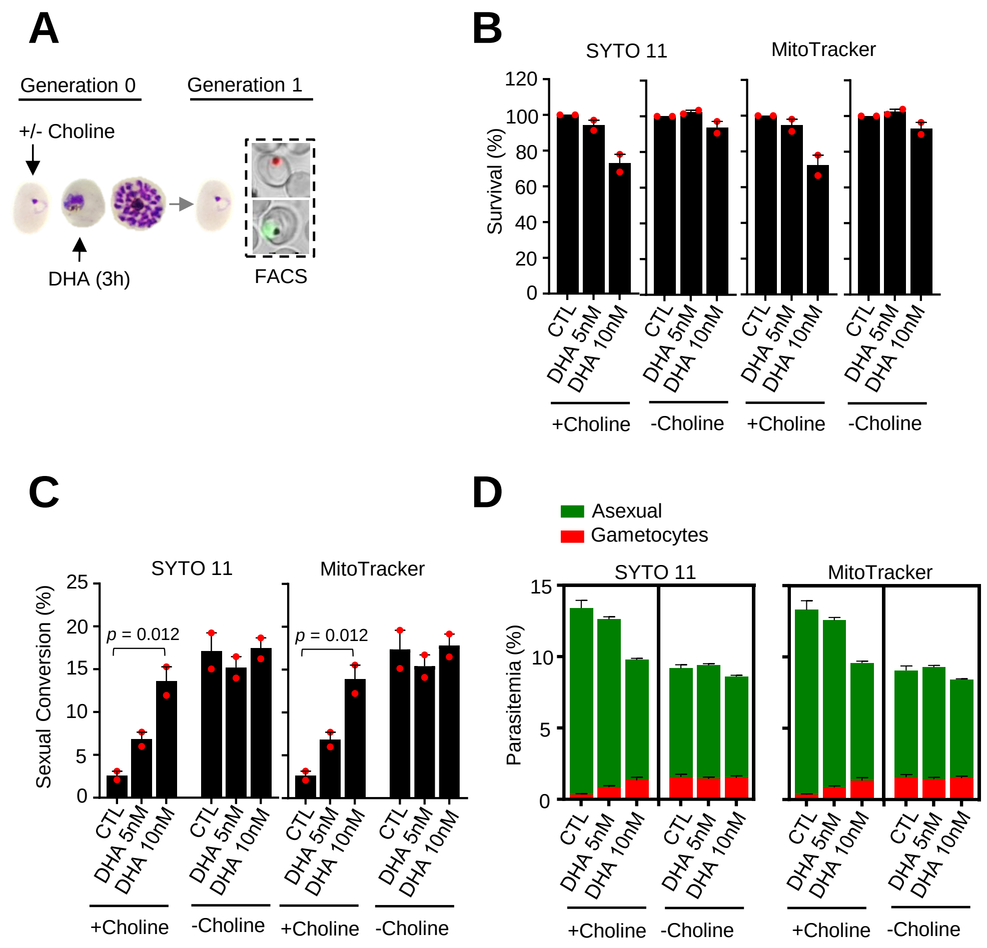


**Supplementary Figure 2. Effect of a dihydroartemisinin (DHA) pulse at the trophozoite stage on sexual conversion in the *E5-gexp02-Tom* line. (A)** Schematic representation of the assay**.** Tightly synchronized cultures of the *E5-gexp02-Tom* line maintained under non-inducing (+ choline) or inducing (- choline) conditions were exposed to a 3 h DHA pulse at subcurative doses at the trophozoite stage (25-30 hpi). Sexual conversion was measured by flow cytometry (FACS) after reinvasion (~30-35 hpi of the next multiplication cycle). **(B)** Survival rate of cultures exposed to the different drug doses, using total parasitemia values (asexual + sexual parasites) based on identification of all parasites or viable parasites only, with SYTO 11 or MitoTracker Deep Red FM, respectively. For each choline condition, values are presented relative to the parasitemia in the control cultures (no drug). **(C)** Sexual conversion rates determined by flow cytometry. The *p* value is indicated only for treatment vs control (no drug) significant differences ($p<0.05$). **(D)** Distribution of absolute parasitemia of asexual and sexual parasites. In all panels, data are presented as the average and s.e.m. of 2 independent biological replicates.

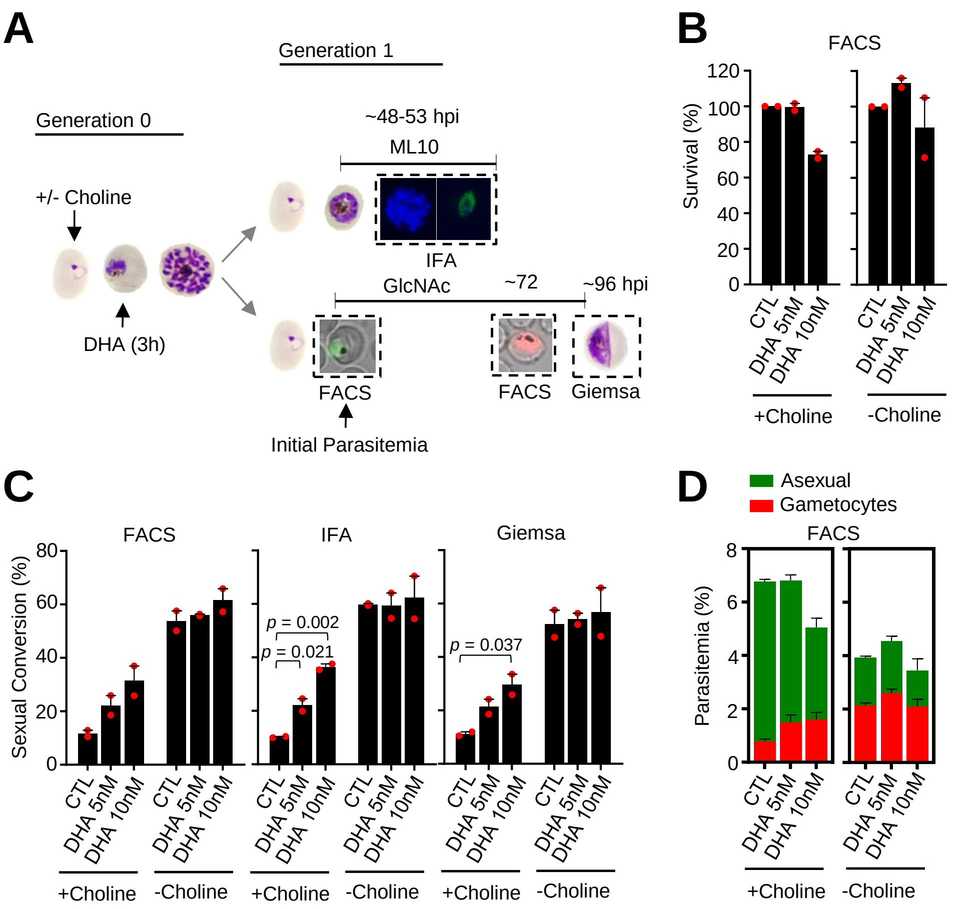


**Supplementary Figure 3. Effect on sexual conversion of a dihydroartemisinin (DHA) pulse at the trophozoite stage in the *NF54-10.3-Tom* line, determined by three different methods. (A)** Schematic representation of the assay. Tightly synchronized cultures of the *NF54-10.3-Tom* line (expression of the fluorescence reporter starts later during gametocyte development than in the *NF54-gexp02-Tom* line) maintained under non-inducing (+ choline) or inducing (- choline) conditions were exposed to a 3 h DHA pulse at subcurative doses at the trophozoite stage (25-30 hpi). Sexual conversion was measured by: (i) flow cytometry analysis (FACS) on D1 (D0 is the first day of Generation 1) to determine the initial parasitemia (SYTO 11), and on D3 to determine the gametocytemia (SYTO 11 and TdTomato), using cultures treated with N-acetylglucosamine (GlcNAc); (ii) immunofluorescence assay (IFA) analysis of cultures treated with ML10 using the Pfs16 marker; (iii) flow cytometry analysis to determine the initial parasitemia, and on D4 microscopy analysis of Giemsa-stained blood smears (Giemsa) to determine the gametocytemia in cultures treated with GlcNAc. **(B)** Survival rate of cultures exposed to the different drug doses, using total parasitemia values (asexual + sexual parasites) determined with the SYTO 11 stain. For each choline condition, values are presented relative to the parasitemia in the control cultures (no drug). **(C)** Sexual conversion rates determined by FACS, IFA, and Giemsa-stained blood smears. The *p* value is indicated only for treatment vs control (no drug) significant differences ($p$<0.05). **(D)** Distribution of absolute parasitemia of asexual and sexual parasites, determined by flow cytometry. In all panels, data are presented as the average and s.e.m. of 2 independent biological replicates.

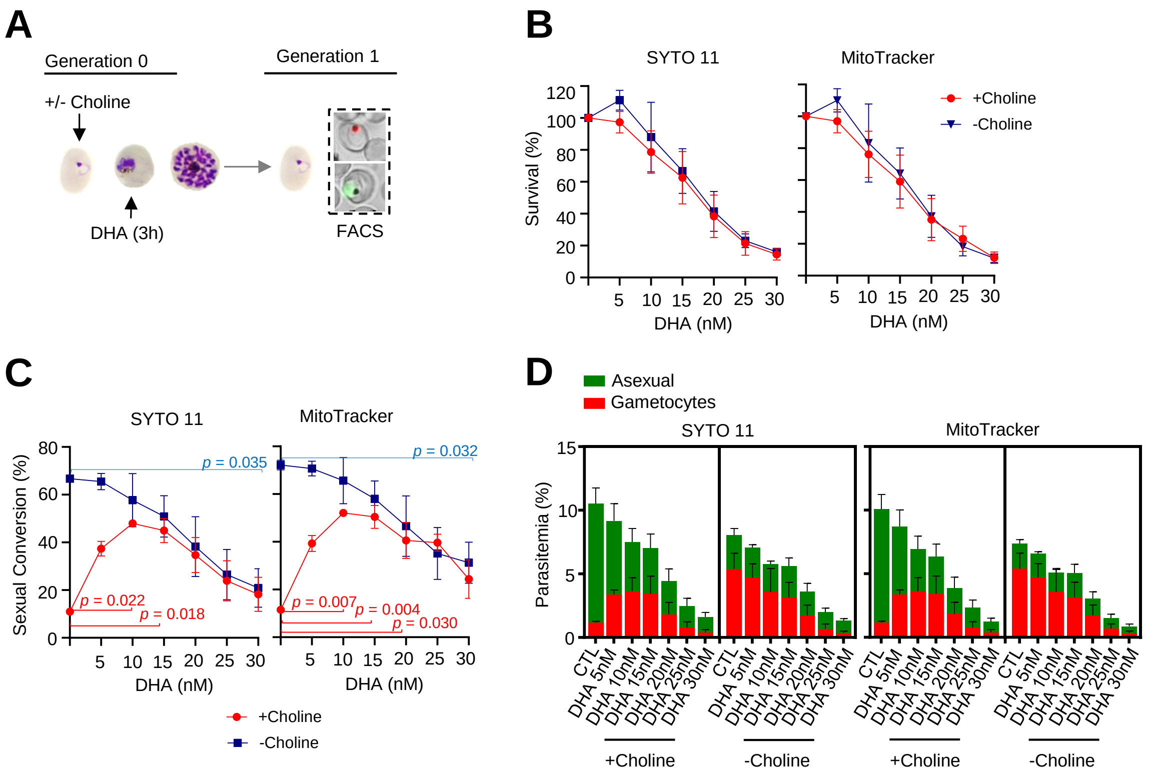


**Supplementary Figure 4. Effect on sexual conversion of a dihydroartemisinin (DHA) pulse at different concentrations during the trophozoite stage. (A)** Schematic representation of the assay. Tightly synchronized cultures of the *NF54-gexp02-Tom* line maintained under non-inducing (+ choline) or inducing (- choline) conditions were exposed to a 3 h DHA pulse at various doses at the trophozoite stage (25-30 hpi). Sexual conversion was measured by flow cytometry (FACS) after reinvasion (~30-35 hpi of the next multiplication cycle). **(B)** Survival rate of cultures exposed to the different drug doses, using total parasitemia values (asexual + sexual parasites) based on identification of all parasites or viable parasites only, with SYTO 11 or MitoTracker Deep Red FM, respectively. For each choline condition, values are presented relative to the parasitemia in the control cultures (no drug). **(C)** Sexual conversion rates determined by flow cytometry. The *p* value is indicated only for treatment vs control (no drug) significant differences ($p<0.05$). **(D)** Distribution of absolute parasitemia of asexual and sexual parasites. In all panels, data are presented as the average and s.e.m. of 3 independent biological replicates.

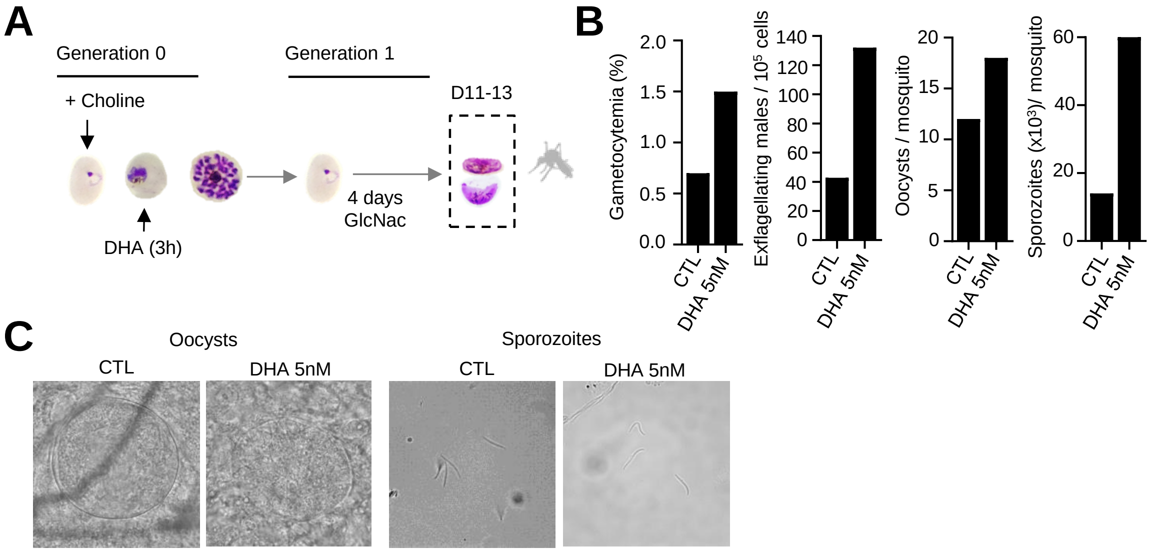


**Supplementary Figure 5. Mosquito infection by gametocytes from cultures exposed to dihydroartemisinin (DHA). (A)** Schematic representation of the assay. Sorbitol-synchronized cultures of the *NF54-gexp02-Tom* line maintained under non-inducing (+ choline) conditions were exposed to a 3 h DHA (5 nM) pulse at the trophozoite stage. On the first day of Generation 1, N-acetylglucosamine (GlcNAc) was added and maintained for four days to eliminate asexual parasites and obtain pure gametocyte cultures. **(B)** Gametocytemia at the time of mosquito infection (11 days after DHA treatment), exflagellation levels (after 10 min of activation with fetal calf serum), mean number of oocysts/mosquito (n=12) and mean number of sporozoites/mosquito (n=10) in a DHA-treated and an untreated control culture (CTL). No morphological differences were observed in gametocyte or mosquito stages between control and DHA-treated cultures. Values are from a single infection experiment and should not be interpreted quantitatively. **(C)** Representative images of oocysts and sporozoites from control and DHA-treated cultures.

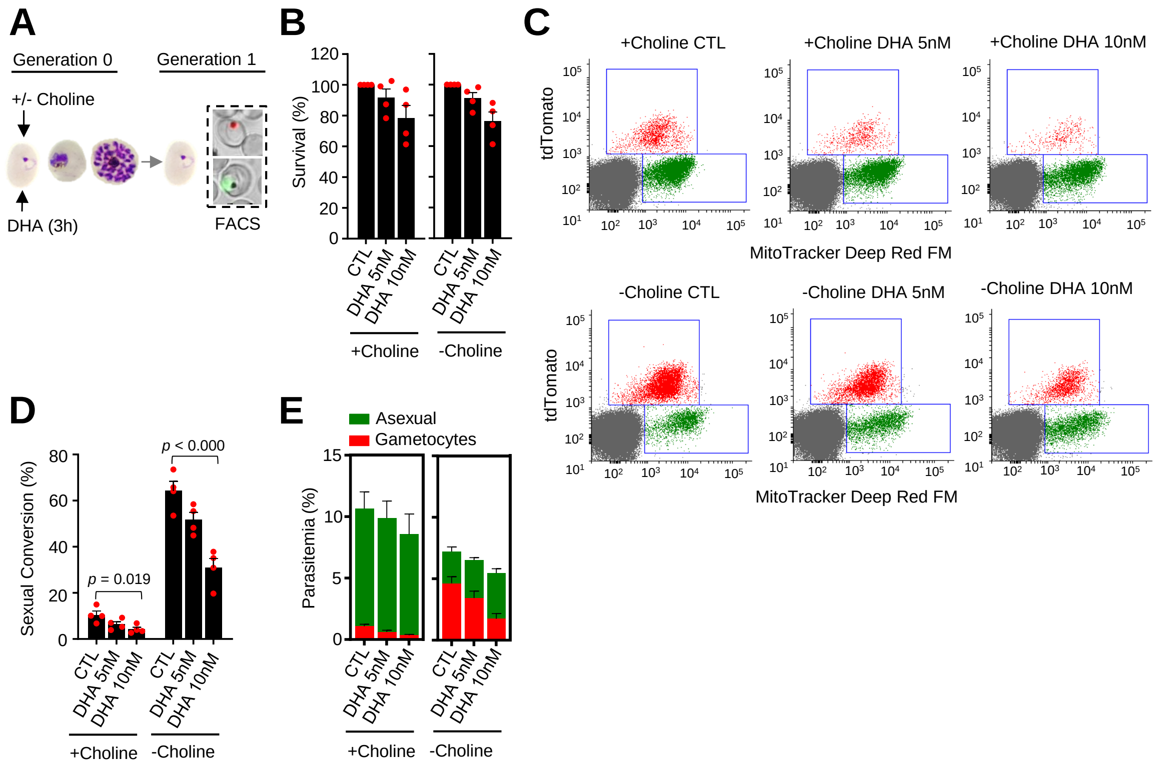


**Supplementary Figure 6. Effect of a dihydroartemisinin (DHA) pulse at the ring stage on sexual conversion, determined using MitoTracker to identify viable parasites. (A)** Schematic representation of the assay. Tightly synchronized cultures of the *NF54-gexp02-Tom* line maintained under non-inducing (+ choline) or inducing (- choline) conditions were exposed to a 3 h DHA pulse at subcurative doses at the early ring stage (0-10 hpi). Sexual conversion was measured by flow cytometry (FACS) after reinvasion (~30-40 hpi of the next multiplication cycle). **(B)** Survival rate of cultures exposed to the different drug doses, using total parasitemia of live parasites (asexual + sexual parasites) determined with a mitochondrial membrane potential stain (MitoTracker Deep Red FM). For each choline condition, values are presented relative to the parasitemia in the control cultures (no drug). **(C)** Representative MitoTracker Deep Red FM vs TdTomato (marks gametocytes) flow cytometry dot plots. **(D)** Sexual conversion rates determined by flow cytometry, calculated using MitoTracker-positive cells only. The *p* value is indicated only for treatment vs control (no drug) significant differences ($p<0.05$). **(E)** Distribution of absolute parasitemia of asexual and sexual parasites. In all panels, data are presented as the average and s.e.m. of 4 independent biological replicates.

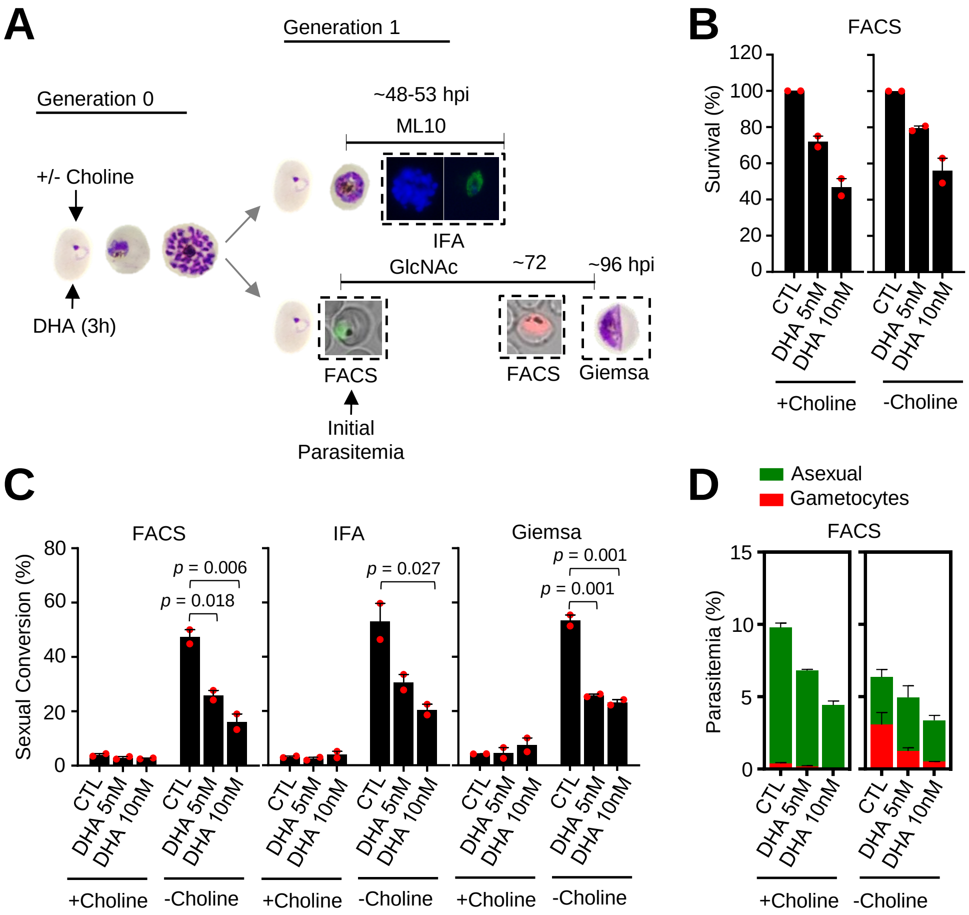


**Supplementary Figure 7. Effect on sexual conversion of a dihydroartemisinin (DHA) pulse at the ring stage in the *NF54-10.3-Tom* line, determined by three different methods. (A)** Schematic representation of the assay. Tightly synchronized cultures of the *NF54-10.3-Tom* line (expression of the fluorescence reporter starts later during gametocyte development than in the *NF54-gexp02-Tom* line) maintained under non-inducing (+ choline) or inducing (- choline) conditions were exposed to a 3 h DHA pulse at subcurative doses at the ring stage (1-6 hpi). Sexual conversion was measured by: (i) flow cytometry analysis (FACS) on D1 (D0 is the first day of Generation 1) to determine the initial parasitemia (SYTO 11), and on D3 to determine the gametocytemia (SYTO 11 and TdTomato), using cultures treated with N-acetylglucosamine (GlcNAc); (ii) immunofluorescence assay (IFA) analysis of cultures treated with ML10 using the Pfs16 marker; (iii) flow cytometry analysis to determine the initial parasitemia, and on D4 microscopy analysis of Giemsa-stained blood smears (Giemsa) to determine the gametocytemia in cultures treated with GlcNAc. **(B)** Survival rate of cultures exposed to different drug doses, using total parasitemia values (asexual + sexual parasites) determined with the SYTO 11 stain. For each choline condition, values are presented relative to the parasitemia in the control cultures (no drug). **(C)** Sexual conversion rates as determined by FACS, IFA, and Giemsa-stained blood smears. The $p$ value is indicated only for treatment vs control (no drug) significant differences ($p<0.05$). **(D)** Distribution of absolute parasitemia of asexual and sexual parasites, determined by flow cytometry. In all panels, data are presented as the average and s.e.m. of 2 independent biological replicates.

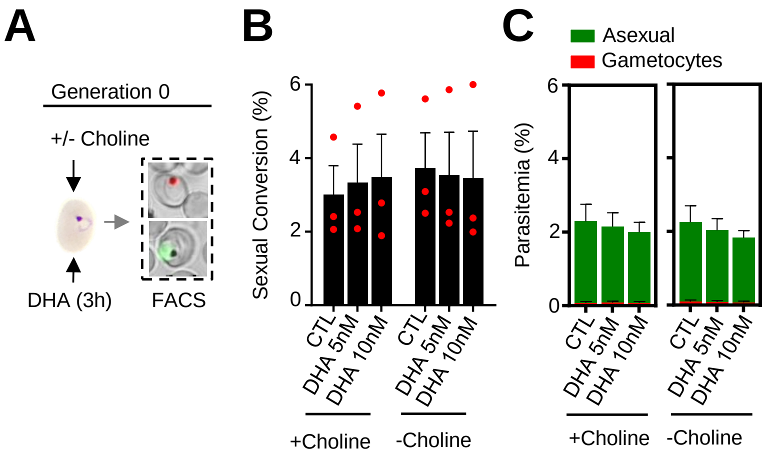


**Supplementary Figure 8. Effect of a dihydroartemisinin (DHA) pulse at the ring stage on sexual conversion by the same cycle conversion (SCC) route, determined using MitoTracker to identify viable parasites. (A)** Schematic representation of the assay. Tightly synchronized cultures of the *NF54-gexp02-Tom* line maintained under non-inducing (+ choline) or inducing (- choline) conditions were exposed to a 3 h DHA pulse at subcurative doses at the early ring stage (0-10 hpi). Sexual conversion was measured by flow cytometry (FACS) within the same cycle (~30-40 hpi) to determine the effect of the drug pulse only on production of new gametocytes by the SSC route. **(B)** Sexual conversion rates determined by flow cytometry, calculated using MitoTracker-positive cells only. No significant difference ($p$<0.05) with the control (no drug) was observed for any treatment condition. **(C)** Distribution of absolute parasitemia of asexual and sexual parasites. In all panels, data are presented as the average and s.e.m. of 3 independent biological replicates.

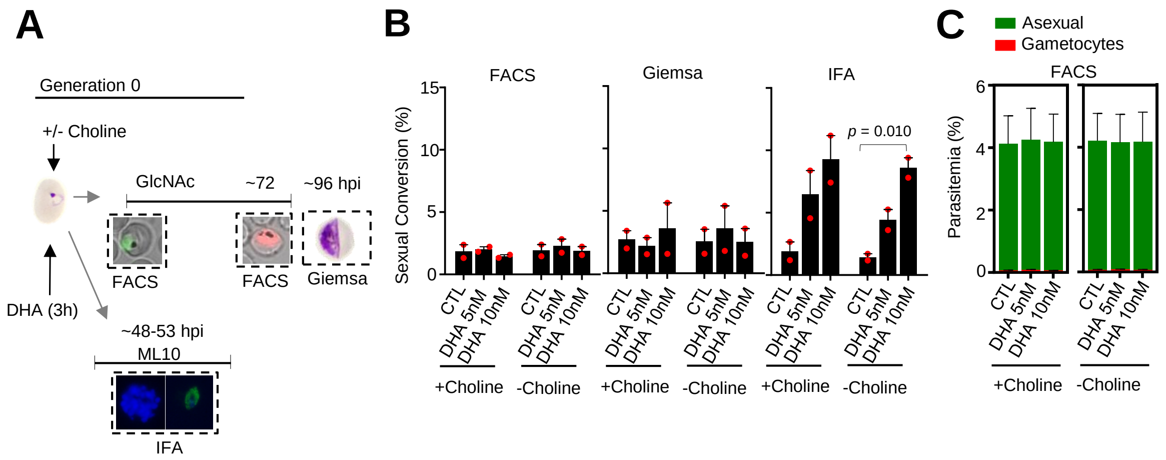


**Supplementary Figure 9. Effect on sexual conversion by the same cycle conversion (SCC) route of a dihydroartemisinin (DHA) pulse at the ring stage in the *NF54-10.3-Tom* line, determined by three different methods. (A)** Schematic representation of the assay. Tightly synchronized cultures of the *NF54-10.3-Tom* line maintained under non-inducing (+ choline) or inducing (- choline) conditions were exposed to a 3 h DHA pulse at subcurative doses at the early ring stage (1-6 hpi). Sexual conversion was measured by: (i) flow cytometry analysis (FACS) on D1 (D0 is the first day of Generation 0, i.e., the day of drug treatment) to determine the initial parasitemia (SYTO 11), and on D3 to determine the gametocytemia (SYTO 11 and TdTomato), using cultures treated with N-acetylglucosamine (GlcNAc); (ii) immunofluorescence assay (IFA) analysis of cultures treated with ML10 using the Pfs16 marker; (iii) flow cytometry analysis to determine the initial parasitemia, and on D4 microscopy analysis of Giemsa-stained blood smears (Giemsa) to determine the gametocytemia in cultures treated with GlcNAc. **(B)** Sexual conversion rates as determined by FACS, IFA, and Giemsa-stained blood smears. The *p* value is indicated only for treatment vs control (no drug) significant differences ($p$<0.05). **(C)** Distribution of absolute parasitemia of asexual and sexual parasites, determined by flow cytometry. In all panels, data are presented as the average and s.e.m. of 2 independent biological replicates.

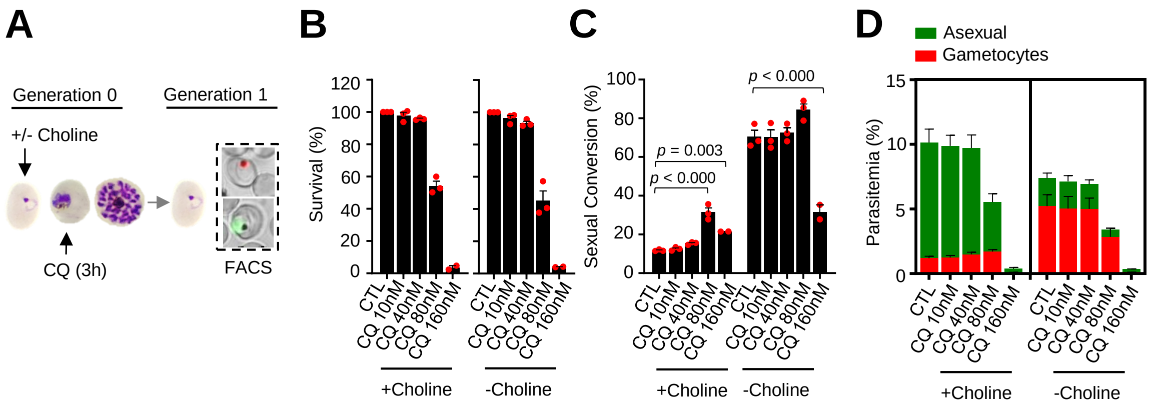


**Supplementary Figure 10. Effect of a chloroquine (CQ) pulse at the trophozoite stage on sexual conversion, determined using MitoTracker to identify viable parasites. (A)** Schematic representation of the assay. Tightly synchronized cultures of the *NF54-gexp02-Tom* line maintained under non-inducing (+ choline) or inducing (- choline) conditions were exposed to a 3 h CQ pulse at subcurative doses at the trophozoite stage (25-30 hpi). Sexual conversion was measured by flow cytometry (FACS) after reinvasion (~30-35 hpi of the next multiplication cycle). **(B)** Survival rate of cultures exposed to the different drug doses, using total parasitemia of live parasites (asexual + sexual parasites) determined with a mitochondrial membrane potential stain (MitoTracker Deep Red FM). For each choline condition, values are presented relative to the parasitemia in the control cultures (no drug). **(C)** Sexual conversion rates determined by flow cytometry, calculated using MitoTracker-positive cells only. The *p* value is indicated only for treatment vs control (no drug) significant differences ($p<0.05$). **(D)** Distribution of absolute parasitemia of asexual and sexual parasites. In all panels, data are presented as the average and s.e.m. of 3 independent biological replicates.

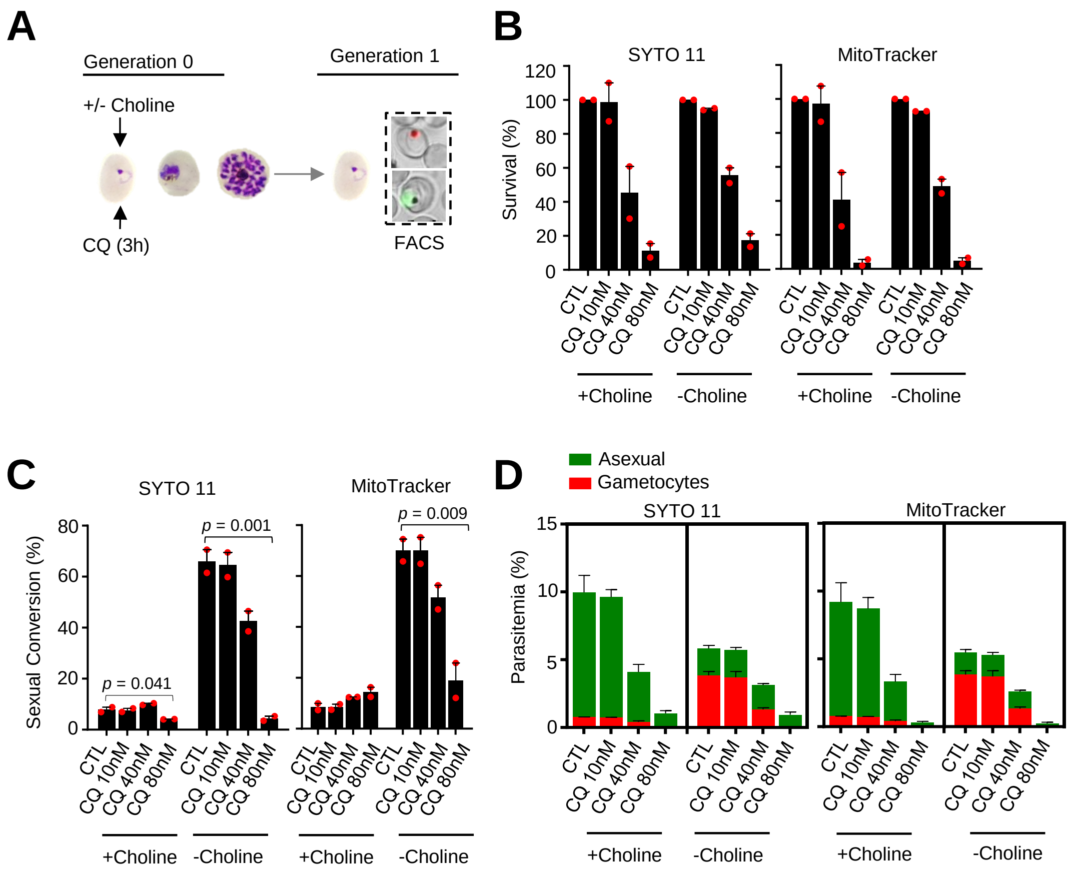


**Supplementary Figure 11. Effect of a chloroquine (CQ) pulse at the ring stage on sexual conversion. (A)** Schematic representation of the assay. Tightly synchronized cultures of the *NF54-gexp02-Tom* line maintained under non-inducing (+ choline) or inducing (- choline) conditions were exposed to a 3 h CQ pulse at subcurative doses at the ring stage (0-10 hpi). Sexual conversion was measured by flow cytometry (FACS) after reinvasion (~30-40 hpi of the next multiplication cycle). **(B)** Survival rate of cultures exposed to the different drug doses, using total parasitemia values (asexual + sexual parasites) based on identification of all parasites or viable parasites only, with SYTO 11 or MitoTracker Deep Red FM, respectively. For each choline condition, values are presented relative to the parasitemia in the control cultures (no drug). **(C)** Sexual conversion rates determined by flow cytometry. The *p* value is indicated only for treatment vs control (no drug) significant differences ($p$<0.05). **(D)** Distribution of absolute parasitemia of asexual and sexual parasites. In all panels, data are presented as the average and s.e.m. of 2 independent biological replicates.

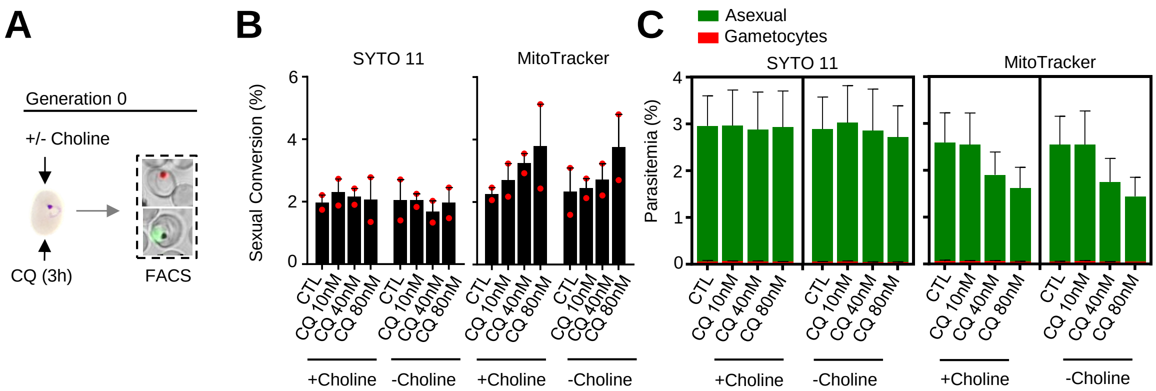


**Supplementary Figure 12. Effect of a chloroquine (CQ) pulse at the ring stage on sexual conversion by the same cycle conversion (SCC) route. (A)** Schematic representation of the assay. Tightly synchronized cultures of the *NF54-gexp02-Tom* line maintained under non-inducing (+ choline) or inducing (- choline) conditions were exposed to a 3 h CQ pulse at subcurative doses at the early ring stage (0-10 hpi). Sexual conversion was measured by flow cytometry (FACS) within the same cycle (~30-40 hpi) to determine the effect of the drug pulse only on production of new gametocytes by the SSC route. **(B)** Sexual conversion rates as determined by flow cytometry using SYTO 11 or MitoTracker Deep Red FM to identify all parasites or viable parasites only, respectively, in addition to TdTomato to identify gametocytes. No significant difference ($p<0.05$) with the control (no drug) was observed for any treatment condition. **(C)** Distribution of absolute parasitemia of asexual and sexual parasites. In all panels, data are presented as the average and s.e.m. of 2 independent biological replicates.

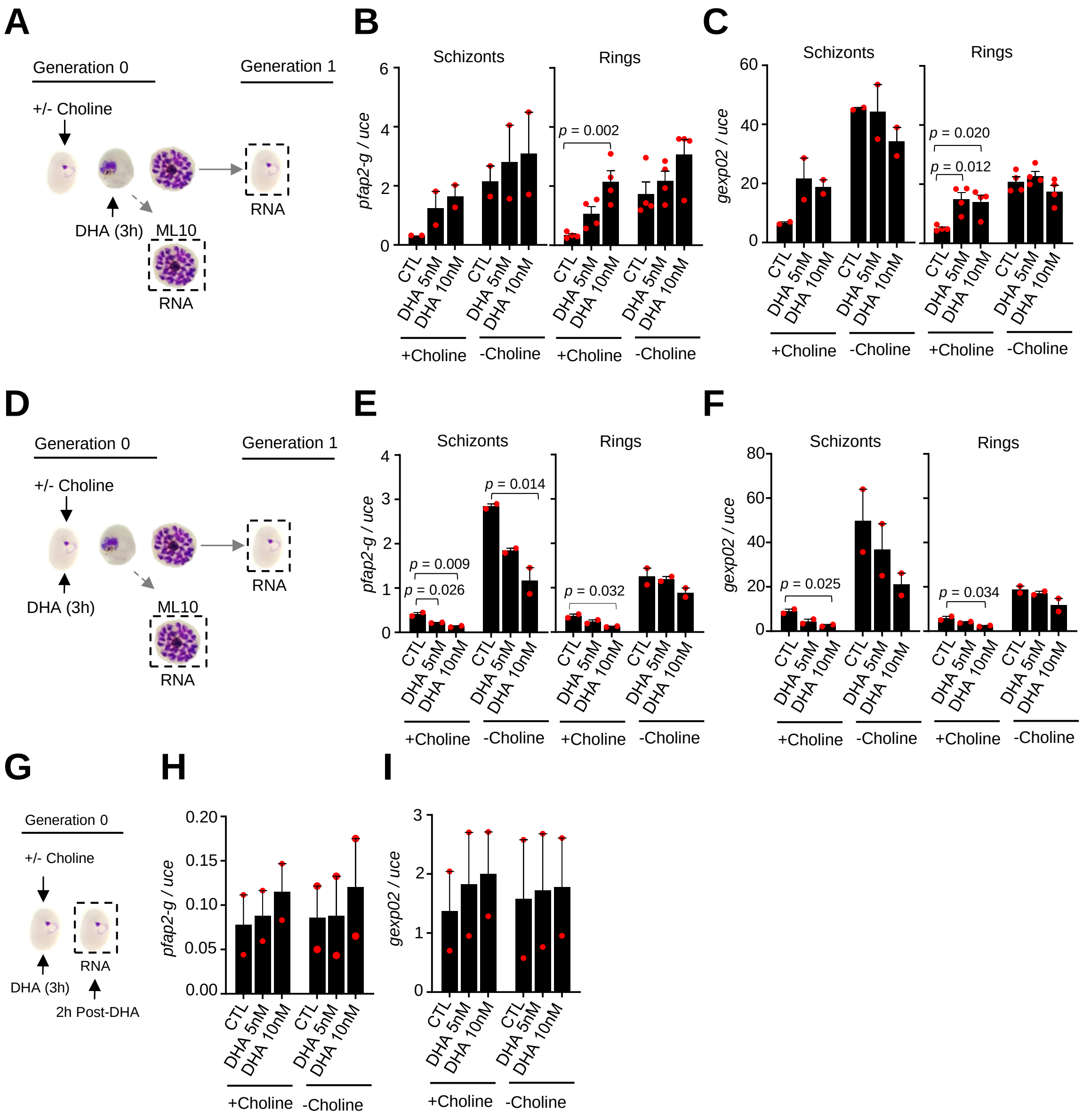

**Supplementary Figure 13. Changes in the expression of *pfap2-g* and *gexp02* after a dihydroartemisinin (DHA) pulse. (A)** Schematic representation of the assay. Tightly synchronized cultures of the *NF54-gexp02-Tom* line maintained under non-inducing (+ choline) or inducing (- choline) conditions were exposed to a 3 h DHA pulse at subcurative doses at the trophozoite stage (25-30 hpi). RNA for transcriptional analysis was collected from ML10-treated cultures at the mature schizont stage (48-53 hpi) and, after reinvasion, from cultures at the early ring stage (cultures not treated with ML10, ~5 hpi). **(B-C)** Transcript levels of *pfap2-g* (B) or *gexp02* (C) normalized against the *ubiquitin-conjugating enzyme* (*uce*) gene. **(D-F)** Same as panels A-C, but cultures were exposed to DHA at the ring stage (0-10 hpi). **(G-I)** Same as panels D-F, but RNA for transcriptional analysis was collected only 2 h after completing the drug pulse. Data are presented as the average and s.e.m. of 4 (panels B-C, rings) or 2 (other panels) independent biological replicates. The *p* value is indicated only for treatment vs control (no drug) significant differences (p<0.05).

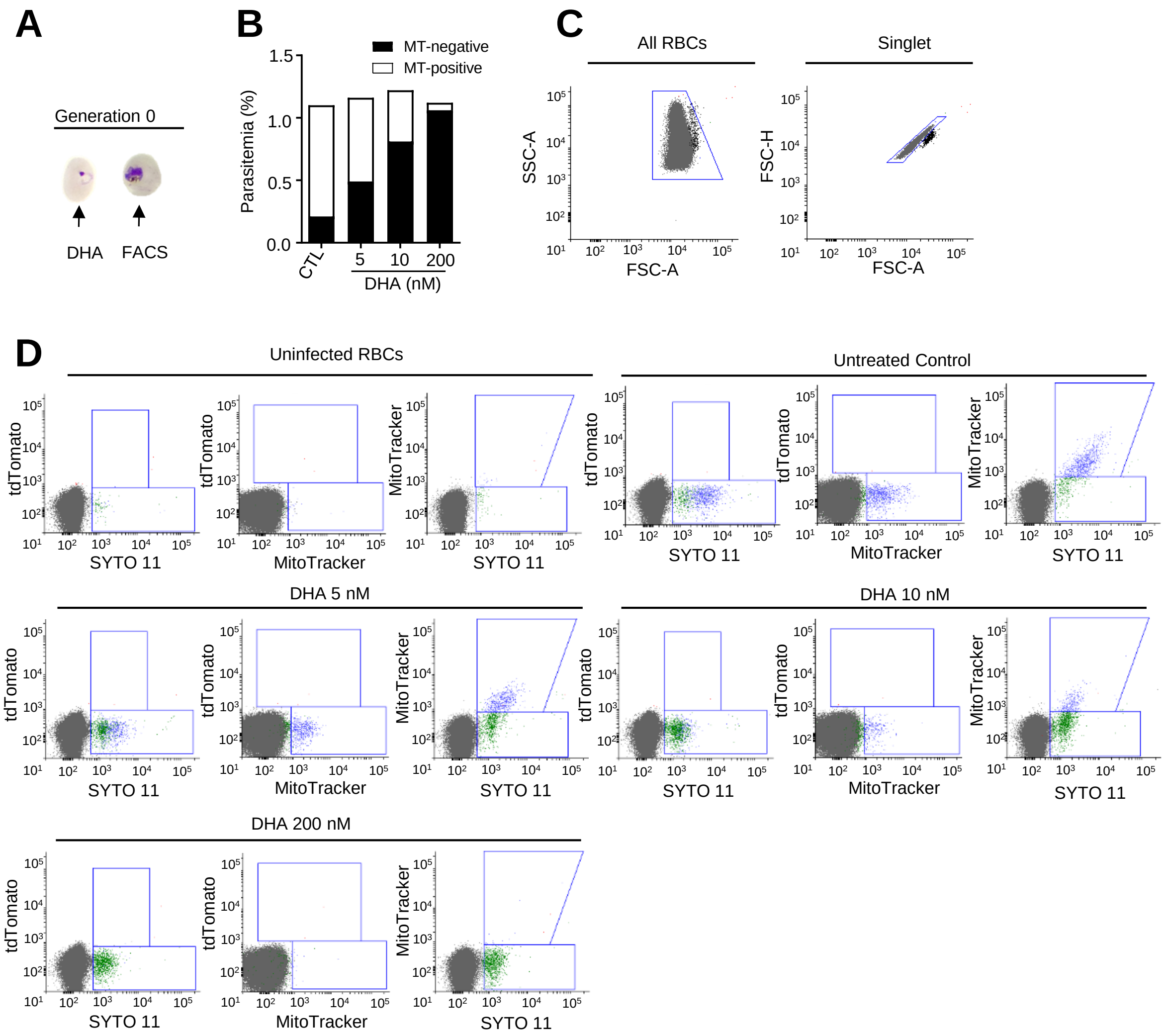


**Supplementary Figure 14. Flow cytometry set-up for the identification of viable parasites in the 3D7-A parasite line using MitoTracker. (A)** Schematic representation of the assay. Tightly synchronized cultures of the 3D7-A wild-type parasite line were exposed to a 3 h dihydroartemisinin (DHA) pulse at subcurative doses at the ring stage (0-20 hpi) or maintained under a lethal dose (200 nM, 'kill' control) for ~24 h. Flow cytometry measurements were performed on the next day, within the same asexual cycle. **(B)** Total parasitemia as determined using SYTO-11, and distribution of MitoTracker (MT) Deep Red FM-positive (viable) and –negative (non-viable and non-stained) parasites after DHA exposure. **(C)** Dot plots of the initial gating strategy. The red blood cell (RBC) population was gated first for cell granularity and size (SSC-A versus FSC-A plot) and then for a defined singlet population (FSC-H versus FSC-A plot). **(D)** Flow cytometry dot plots for MitoTracker Deep Red FM, SYTO 11, and TdTomato (marks gametocytes in the transgenic lines, absent in wild type parasites). Some presumably healthy parasites were not stained with MitoTracker, as revealed by the presence of MitoTracker-negative/SYTO 11-positive parasites in the no-drug control.

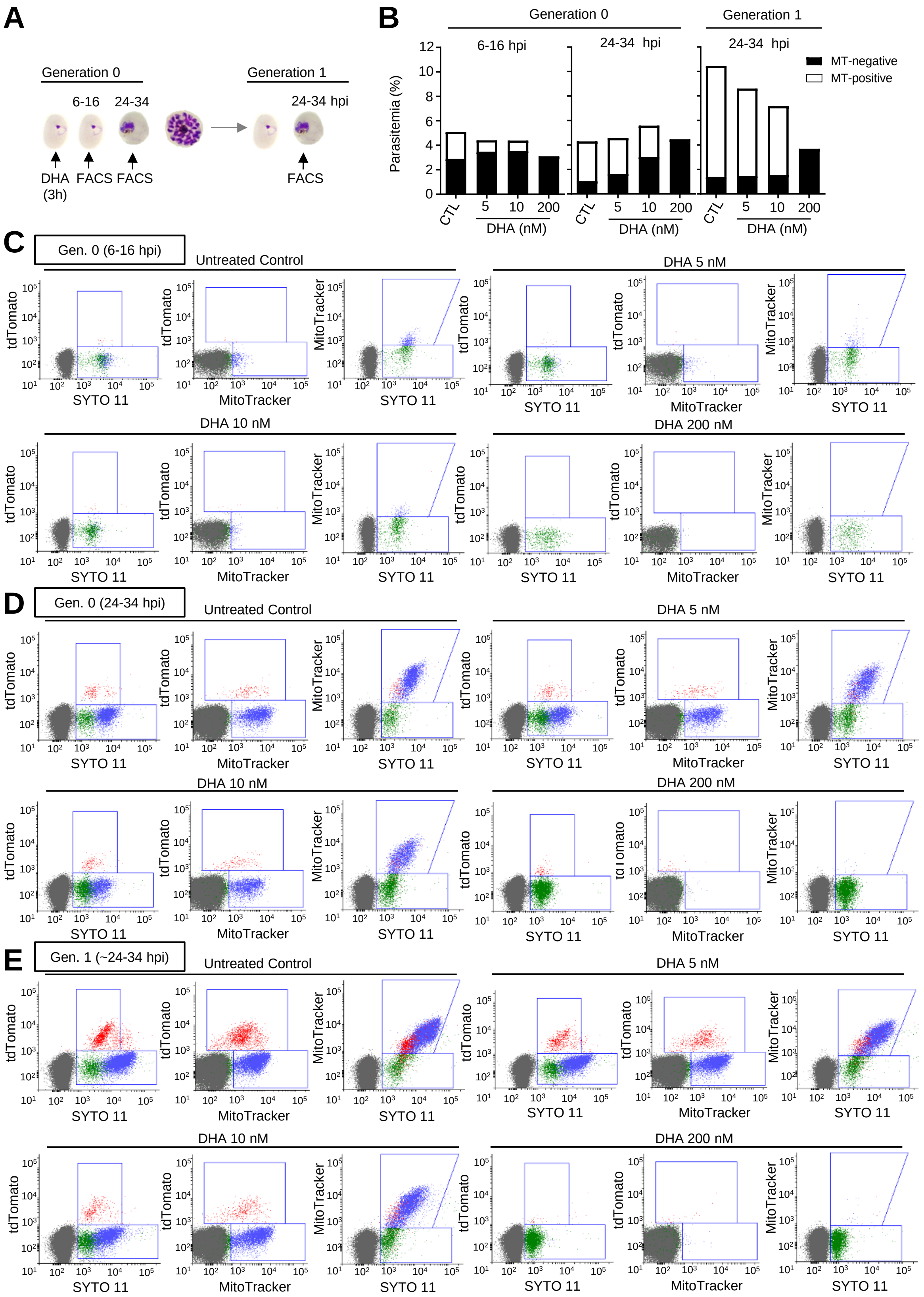


**Supplementary Figure 15. Flow cytometry set-up for the identification of viable parasites in the *NF54-gexp02-Tom* line using MitoTracker. (A)**

Schematic representation of the assay. Tightly synchronized cultures of the *NF54-gexp02-Tom* line were exposed to a 3 h dihydroartemisinin (DHA) pulse at subcurative doses at the ring stage (0-10 hpi) or maintained under a lethal dose (200 nM, ‘kill’ control) for up to 48 h. Flow cytometry measurements were performed at different times within the same cycle, and after reinvasion, as indicated. **(B)** Total parasitemia as determined using SYTO-11, and distribution of MitoTracker (MT) Deep Red FM-positive (viable) and –negative (non-viable and non-stained) parasites after DHA exposure. **(C-E)** Flow cytometry dot plots for MitoTracker Deep Red FM, SYTO 11, and TdTomato (marks gametocytes) at 6-16 hpi (C) and 24-34 hpi (D) of the same cycle of DHA treatment, and ~24-34 hpi of the next cycle (E). Some presumably healthy parasites were not stained with MitoTracker, as revealed by the presence of MitoTracker-negative/SYTO 11-positive parasites in the no-drug control (especially in ring-stage cultures).
